# Macrophage Released Pyrimidines Inhibit Gemcitabine Therapy in Pancreatic Cancer

**DOI:** 10.1101/463125

**Authors:** Christopher J. Halbrook, Corbin Pontious, Ho-Joon Lee, Ilya Kovalenko, Yaqing Zhang, Laura Lapienyte, Stephan Dreyer, Daniel M. Kremer, Peter Sajjakulnukit, Li Zhang, Barbara Nelson, Hanna S. Hong, Samantha Kemp, David Chang, Andrew Biankin, Howard C. Crawford, Jennifer P. Morton, Marina Pasca di Magliano, Costas A. Lyssiotis

## Abstract

Pancreatic Ductal Adenocarcinoma (PDA) is characterized by abundant infiltration of tumor associated macrophages (TAMs). TAMs have been reported to drive resistance to gemcitabine, the front-line chemotherapy in PDA, though the mechanism of this resistance remains unclear. Profiling metabolite exchange, we demonstrate macrophages programmed by PDA cells release a spectrum of pyrimidine species. These include deoxycytidine, which inhibits gemcitabine through molecular competition at the level of drug uptake and metabolism. Accordingly, genetic or pharmacological depletion of TAMs in murine models of PDA sensitizes these tumors to gemcitabine. Consistent with this, patients with low macrophage burden demonstrate superior response to gemcitabine treatment. Additionally, we report pyrimidine release is a general function of anti-inflammatory myeloid cells, suggesting an unknown physiological role of pyrimidine exchange by immune cells.

## Introduction

Pancreatic Ductal Adenocarcinoma (PDA), one of the deadliest human cancers (Siegel et al., 2016), is characterized by a dense matrix rich with activated fibroblasts and tumor-associated macrophages (TAMs). Patients respond poorly to current treatments, and the degree of therapeutic resistance correlates with the fibroinflammatory response (Koay et al., 2014). This reaction inhibits vascularization and/or vessel function, and thus presumably delivery of therapeutic agents (Feig et al., 2012; Olive et al., 2009; Provenzano et al., 2012; Rhim et al., 2014). Accordingly, nutrient acquisition and metabolism pathways are rewired in PDA cells to support survival and growth in this hypoxic, avascular tumor microenvironment (Halbrook and Lyssiotis, 2017; Perera and Bardeesy, 2015). For example, we recently demonstrated that PDA cells obtain nutrients through metabolic crosstalk with cancer-associated fibroblasts (CAFs) (Sousa et al., 2016), a cell type in PDA tumors that can outnumber malignant cells by as much as ten to one. In addition to CAFs, TAMs constitute a large proportion of the overall cellularity and are important regulators of the tumor microenvironment (Di Caro et al., 2016). TAM abundance correlates with a worse prognosis in PDA (Di Caro et al., 2016), systemic TAM depletion can block pancreatic tumorigenesis and regress established PDA tumors (Mitchem et al., 2013; Zhang et al., 2017a), and have been suggested to inhibit chemotherapy through unknown means (Mitchem et al., 2013; Weizman et al., 2014).

In physiological settings, inflammatory and anti-inflammatory properties of macrophages can be directed and mediated by cellular metabolism programs (Van den Bossche et al., 2017).

Similarly, TAM functions are also shaped by cell intrinsic metabolism and the functional consequences of this metabolism on the tumor microenvironment (Lyssiotis and Kimmelman, 2017; Murray, 2016). Based on this, and the abundance of TAMs in PDA tumors, we hypothesized that TAMs may influence therapeutic response in PDA tumors through metabolic crosstalk with cancer cells.

## Results

### Macrophage released pyrimidines confer gemcitabine resistance to PDA cells

To study metabolite crosstalk between macrophages and PDA cells, we generated tumor educated macrophages (TEMs) by polarizing murine bone marrow-derived macrophages (BMDMs) with PDA cell conditioned media (CM) (Fig. S1a) using a system previously established to mimic pancreatic TAMs in vivo (Zhang et al., 2017a; Zhang et al., 2017b). In parallel, BMDMs were directed to a classically activated phenotype (i.e. M1) by treatment with lipopolysaccharide (LPS) or polarized to an alternatively activated phenotype (i.e. M2) by treatment with Interleukin 4 (IL4) (Van den Bossche et al., 2017). After 48 hours, conditioned media from these cultures were profiled using liquid chromatography-coupled tandem mass spectrometry (LC-MS/MS) metabolomics (Fig. 1a). This analysis revealed accumulation of pyrimidine nucleosides and nucleobases in TEM media (Fig. 1b). In contrast, we did not observe a similar profile for release of purine species. When PDA cells were then incubated in TEM CM, many of the accumulated pyrimidines were depleted from TEM media by PDA cells (Fig. 1c), suggesting a directional transfer of these metabolites. This observation was provocative for several reasons. First, pyrimidine release and their transfer among cells has neither been previously reported nor characterized. Further, the front-line chemotherapy for PDA is gemcitabine (Gem), a pyrimidine anti-nucleoside. Gem resistance has been linked to TAMs in PDA (Mitchem et al., 2013), although the mechanism behind this link remains unclear. Thus, we hypothesized that pyrimidine nucleosides released by TEMs may directly confer gem resistance to PDA cells.

**Figure 1:**
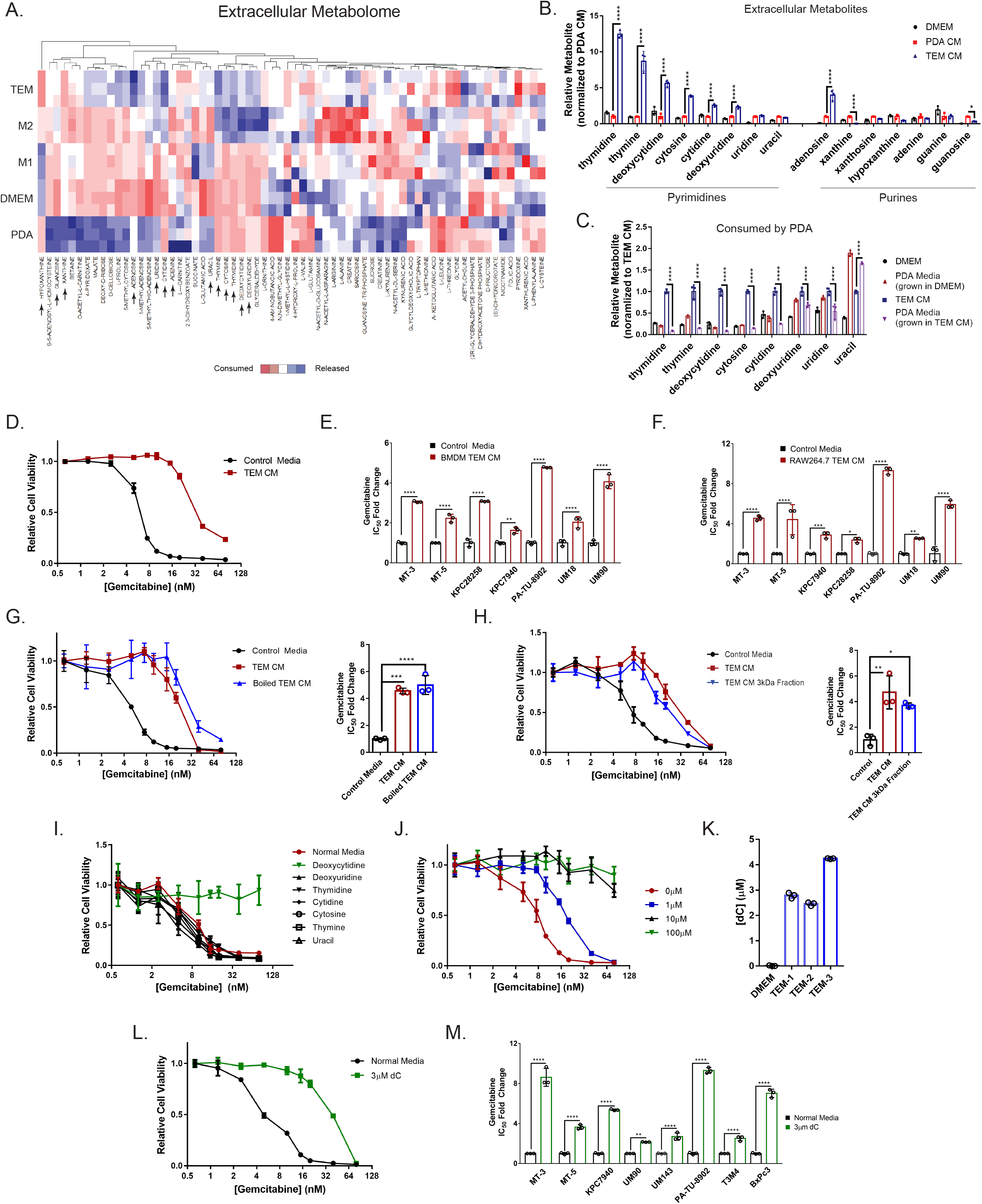
Tumor educated macrophages confer gemcitabine resistance to PDA cells through deoxycytidine release. **1a**. Heat map of metabolites in the conditioned media (CM) of tumor educated macrophages (TEM), alternatively activated macrophages (M2), classically-activated macrophages (M1), iKras*3 PDA cell line, and complete DMEM. Blue represents relative metabolite release, red represents relative metabolite consumption. Metabolites noted with an arrow are presented in the bar graph in (1b). Macrophage subtypes are polarized from matched biological replicates (n=3). **1b.** Relative quantification of nucleoside species found in complete DMEM, PDA CM, and TEM CM, normalized to PDA CM (n=3). **1c**. Relative quantification of pyrimidine nucleosides remaining in PDA CM, TEM CM, or DMEM after 24 hours of culture with iKras*3 PDA cells, normalized to TEM CM (n=3). **1d**. Relative viability dose response curve of MT3-KPC cells treated with gemcitabine in the presence of 75% TEM CM vs. control media (n=3) **1e**. Relative fold change of gemcitabine IC_50_ between control or 75% CM from bone marrow-derived macrophages (BMDM) polarized to TEMs or **1f**. RAW 264.7 macrophages polarized to TEMs (n=3). **1g**. Relative viability dose response curve and IC_50_ of MT3-KPC cells treated with gemcitabine in the presence of 75% TEM CM, heat denatured TEM CM, or control media (n=3). **1h**. Relative viability dose response curve and IC_50_ of MT3-KPC cells treated with gemcitabine in the presence of 75% TEM CM, 75% TEM CM passed through a 3 kDa filter, or control media (n=3). **1i**. Relative viability dose response curve of MT3-KPC cells treated with gemcitabine in the presence of 100μM of the indicated pyrimidine in complete DMEM or normal complete DMEM alone (n=3). **1j**. Relative viability dose response curve of MT3-KPC cells treated with gemcitabine in the presence of the indicated concentration of deoxycytidine (dC) in complete DMEM (n=3). **1k**. Calculated abundance of dC from TEM CM generated by 3 independent TEM preparations, compared to base media, determined via LC-MS/MS (n=3). **1l**. Relative viability dose response curve of MT3-KPC cells treated with gemcitabine in the presence of 3μM dC vs. normal complete DMEM (n=3). **1m.** Relative fold change of gemcitabine IC_50_ for cells treated with 3μM dC versus normal complete DMEM (n=3). * *P* ≤ 0.05; ^**^ *P* ≤ 0.01; ^***^ *P* ≤ 0.001; ^****^ *P* ≤ 0.0001.

Accordingly, we ran gem dose response curves in the presence of TEM CM (Fig. 1d). We observed a shift in gem sensitivity with TEM CM, and this result was reproducible across a large panel of cell lines, including primary human patient-derived cultures (Fig. 1e, Fig. S1b). This observation was also true of TEMs generated from a murine macrophage cell line (Fig. 1f, Fig S1c). We further that observed TEM CM retained the ability to confer gem resistance after boiling or passage through a 3kDa cutoff filter (Fig. 1 g,h), providing evidence for a metabolite factor driving the chemoresistance phenotype.

To determine if pyrimidines found in TEM media confer gem resistance, we treated PDA cells with gem in the presence of a high concentration (100μM) of these nucleosides (Fig. 1i). Among these, we found deoxycytidine (dC) uniquely blocked cytotoxic activity of gem in PDA cells, and, furthermore, that this response was dose dependent (Fig. 1j). We then quantitated dC and other pyrimidines in TEM CM by mass spectrometry and observed them to be in the micromolar range (Fig. 1k, Table S1). Importantly, we were able to phenocopy the gem-resistance activity of TEM CM by simply supplementing growth media with 3 μM dC (Fig. 1l, m, Fig. S1d), the concentration observed in TEM CM.

### Pyrimidine release is a property of alternatively activated macrophage metabolism

In addition to TEMs, M2 macrophages also released dC (Fig. 1a), and their media conferred gem resistance to PDA cells (Fig. 2a). Neither feature was observed for M1 macrophage media. The release of pyrimidines from TEMs and M2 macrophages is a biosynthetic and bioenergetically intensive process. And, given the dichotomy between macrophage subtypes, we were interested in understanding how the metabolic programs were wired to facilitate pyrimidine release. The functional metabolism of classical macrophage subtypes and their preferred bioenergetic pathways has been previously characterized (Jha et al., 2015). Anti-inflammatory macrophages were reported to preferentially utilize mitochondrial respiration and fatty acid oxidation (FAO) (Vats et al., 2006), which we also observed in pancreatic TEMs (Fig. 2b-e, Fig. S2a-c).

**Figure 2:**
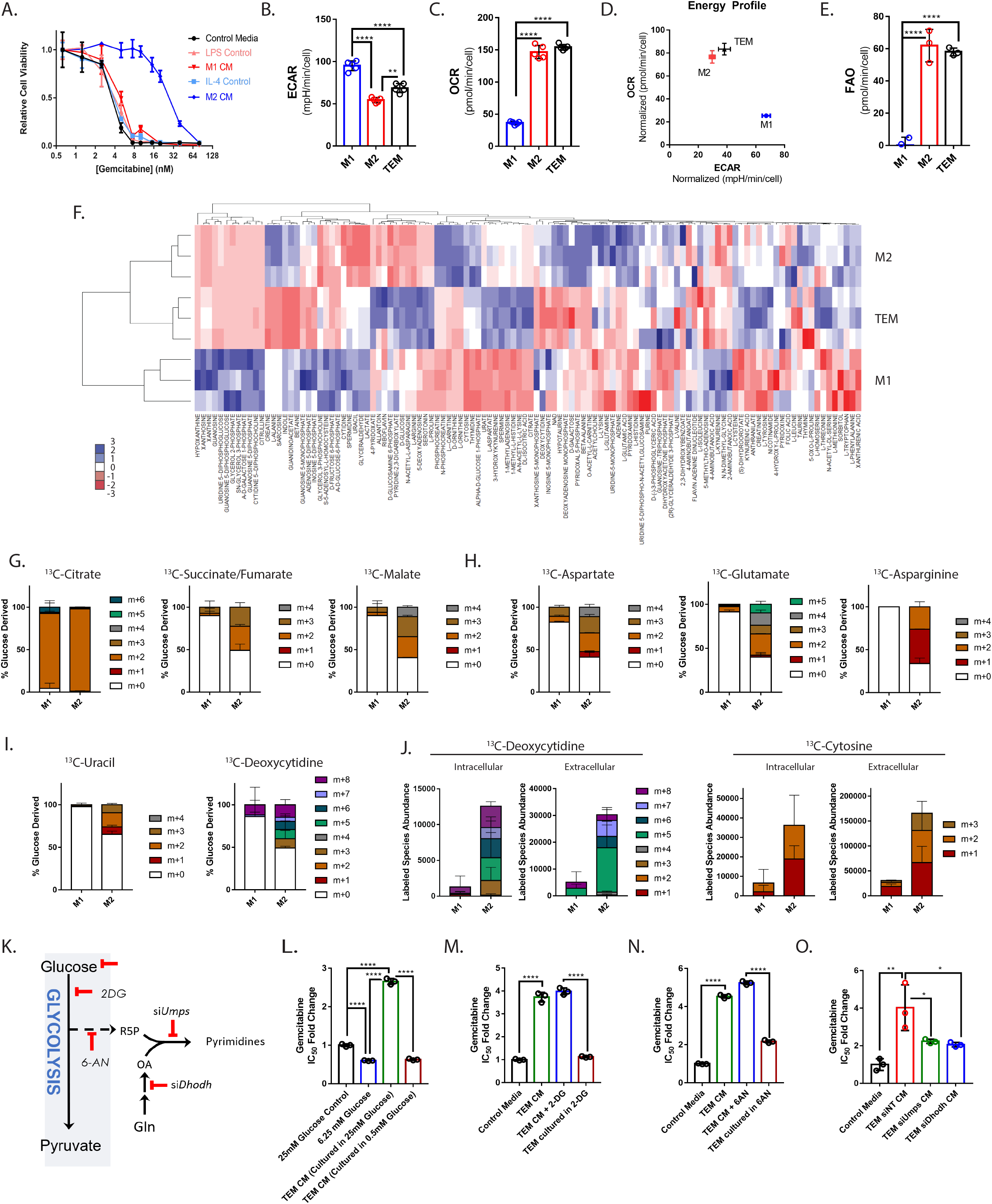
Oxidative metabolism of TEMs and M2 macrophages promote pyrimidine biosynthesis from glucose. **2a**. Relative viability dose response curve of MT3-KPC cells treated with gemcitabine in the presence of M1 or M2 conditioned media (CM) vs. control media (n=3). **2b**. Representative basal extracellular acidification rates (ECAR) and **2c**. oxygen consumption rates (OCR) of TEM, M1, and M2 macrophages (n=5). **2d**. Comparative energy profile of TEM, M1, and M2 macrophages comparing ECAR vs. OCR. **2e**. Basal rate of exogenous fatty acid oxidation (FAO) of TEM, M1, and M2 macrophages (n=5). **2f**. Heatmap representation of significant intracellular metabolites found in TEM, M1, and M2 macrophages by metabolomics profiling (n=3). **2g**. Fractional isotopologue labeling of TCA cycle metabolites, **2h.** amino acids, and **2i.** pyrimidines after 16 hours of labeling with uniformly ^13^C-glucose in M1 vs. M2 macrophages (n=3). **2j**. Intra and extracellular abundance of pyrimidine isotopologues labeled as in (**2g-i**) (n=3). **2k**. Simplified pyrimidine biosynthesis pathway diagram. 2-DG, 2-deoxyglucose; 6-AN, 6-aminonicotinamide; Dhodh, Dihydroorotate dehydrogenase; Gln, glutamine; R5P, ribose 5-phosphate; OA, Orotate; Umps, Uridine 5’-monophosphate synthase. **2l**. Relative fold change in the calculated gemcitabine IC_50_ in KPC-MT3 cells treated in normal glucose DMEM (25mM), 75% glucose restricted media (6.25mM), 75% CM from TEMs grown in normal glucose DMEM, or 75% CM from TEMs grown in glucose restricted media (n=3). **2m**. Relative fold change in the calculated gemcitabine IC_50_ in KPC-MT3 cells treated in normal DMEM, normal DMEM + 200μM 2-deoxyglucose (2-DG), 75% TEM CM, or 75% CM from TEMs grown in 200μM 2-DG (n=3). **2n**. Relative fold change in the calculated gemcitabine IC_50_ in KPC-MT3 cells treated in normal DMEM, normal DMEM + 1μM 6-aminonicotinamide (6-AN), 75% TEM CM, or 75% CM from TEMs grown in 1μM 6-AN (n=3). **2o**. Relative fold change in the calculated gemcitabine IC_50_ in KPC-MT3 cells treated in normal DMEM, or 75% CM from TEMs transfected with siRNA targeting Dhodh, Umps, or nontargeting (NT) siRNA (n=3). * *P* ≤ 0.05; ^**^ *P* ≤ 0.01; ^****^ *P* ≤ 0.0001

These results led us to examine macrophage intracellular metabolomic profiles, which showed a close but not complete correlation between M2 macrophages and TEMs (Fig. 2f). Pathway enrichment analysis of differential metabolites indicated that nucleoside metabolism pathways were highly represented among those that distinguish TEMs and M2 macrophages from M1 macrophages (Table S2). To gain further insight, we carried out stable isotope tracing using uniformly labeled ^13^C-Glucose. The necessity of using PDA conditioned media to polarize TEMs prevented sufficient label incorporation into TEM biosynthetic pathways, limiting our tracing analysis to M1 and M2 macrophages. Nevertheless, comparison between M1 and M2 macrophage metabolism was illuminating, and subsequently relevant in TEMs. Consistent with our bioenergetic profiling analysis (Fig. 2b-e), ^13^C-Glucose fractional labeling patterns revealed increased glucose flux through the tricarboxylic acid (TCA) cycle in M2 macrophages (Fig. 2g), which facilitated TCA cycle anaplerosis (Fig. 2h) and ultimately pyrimidine biosynthesis (Fig. 2i, Fig. S2e). Furthermore, M2 macrophages demonstrated an increased abundance of intercellular and extracellular de novo synthesized pyrimidines as compared to M1 macrophages (Fig. 2j), supporting the idea that pyrimidine release is a property of anti-inflammatory macrophages. Overall, M2 macrophages exhibited a vastly increased biosynthetic capacity from glucose, relative to M1 macrophages (Fig. S2f).

We next sought to determine if targeting pyrimidine production by inhibiting nodes of glucose metabolism would impair the ability of TEMs to confer gem resistance (Fig. 2k). Limiting availability of glucose, inhibiting glycolysis with 2-deoxyglucose (2-DG), or blocking glucose incorporation into the oxidative arm of the pentose phosphate pathway with 6-aminonicotinamide (6-AN) all inhibited the ability of TEM CM to modulate gem sensitivity in PDA cells (Fig. 2l-n, Fig. S3a-c). Importantly, these treatments had a minimal impact on TEM proliferative capacity (Fig. S3d). Similar results were obtained by directly disrupting key pyrimidine biosynthesis enzymes, *Dhodh* and *Umps*, using a genetic knockdown strategy (Fig. 2o, Fig. S3e, f). These results demonstrate the requirement of de novo pyrimidine biosynthesis for TEM CM to inhibit gem activity in PDA cells.

### Deoxycytidine competitively inhibits gemcitabine uptake and metabolism

Gem is a prodrug that requires activation through phosphorylation before it can be inserted into DNA. The first step in this process, phosphorylation by deoxycytidine kinase (dCK), serves to trap gem inside the cell. Subsequently, monophosphorylated-gem undergoes di-, and tri-phosphorylation before incorporation into DNA, where it causes masked chain termination and ultimately cell death (Fig. 3a) (Parker, 2009). Given the structural similarity between dC and gem, we hypothesized that dC could act to inhibit gem activity through molecular competition. To examine this, we measured levels of intra and extracellular gem in the presence or absence of dC and observed that dC treatment resulted in an accumulation of gem both inside and outside the cell (Fig. 3b, Fig S3g). The intercellular accumulation suggested a lack of gem processing and incorporation into DNA. To test this, we further treated PDA cells with ^3^H-radiolabeled gem in the presence or absence of dC and observed that dC treatment prevented gem incorporation into DNA (Fig. 3c). These results suggest that dC is acting to decrease the effective concentration of gem and its activated species experienced by the cell.

**Figure 3:**
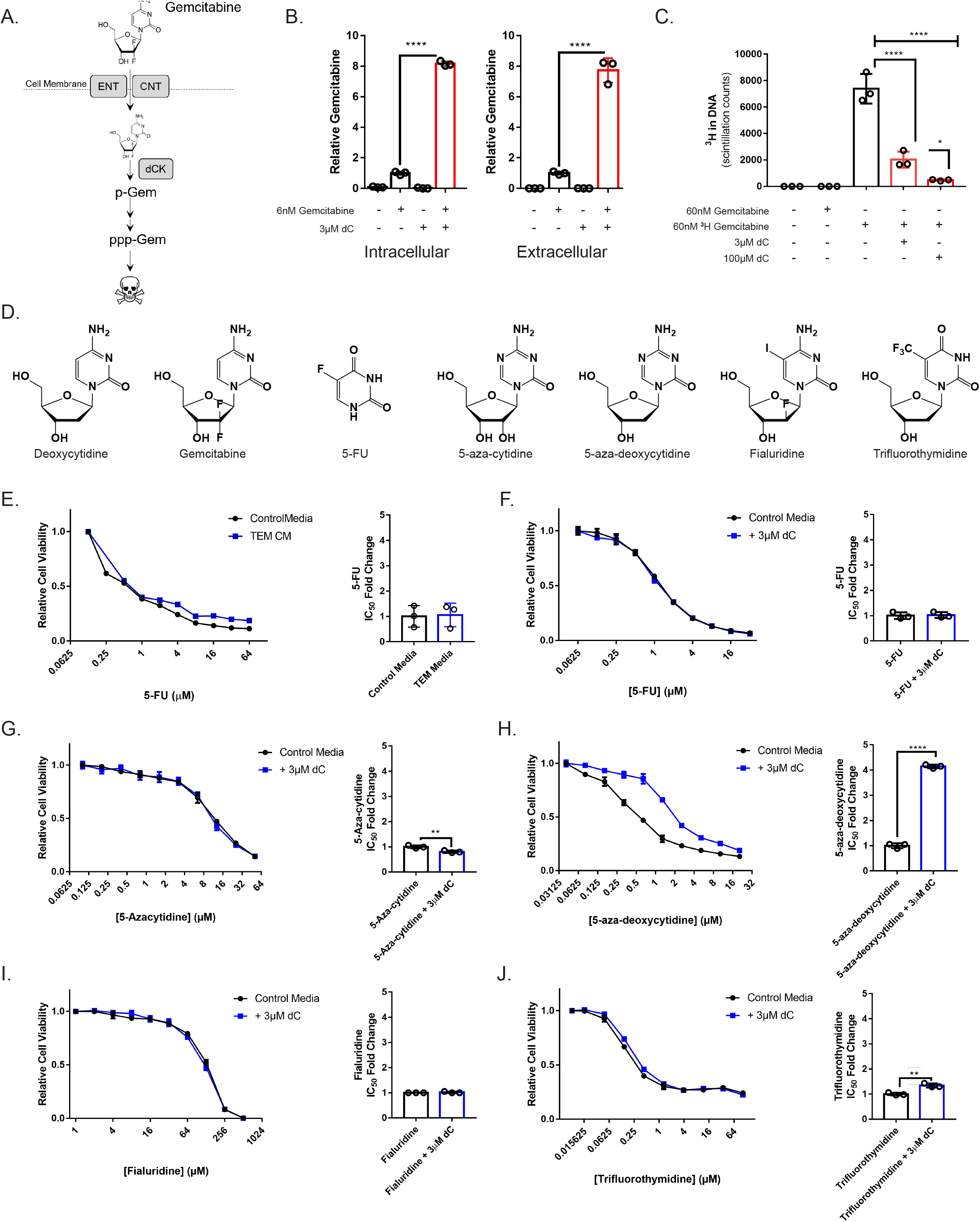
Deoxycytidine blocks the uptake and incorporation of gemcitabine and other dCK activated pyrimidine-based chemotherapies. **3a**. Schematic representation of the mechanism of gemcitabine uptake and metabolism. Gem, gemcitabine; ENT, Equilibrative nucleoside transporter; CNT, Concentrative nucleoside transporter; dCK, deoxycytidine kinase; p-Gem, gemcitabine monophosphate; ppp-gem, gemcitabine triphosphate. **3b**. Relative intra and extracellular abundance of gemcitabine in KPC-MT3 cells or conditioned media, respectively, after 16 hours of treatment with 6nM gemcitabine in the presence or absence of 3μM deoxycytidine, as measured by LC-MS/MS (n=3). **3c**. Incorporation of gemcitabine into the DNA in KPC-MT3 cells treated with 60nM ^3^H-labeled gemcitabine in the presence or absence of the indicated amounts of deoxycytidine (n=3). **3d**. Chemical structures of deoxycytidine and pyrimidine antimetabolite chemotherapies. **3e**. Relative viability dose response curve and IC_50_ of MT3-KPC cells treated with 5-FU in the presence of 75% TEM conditioned media vs. control media or **3f**. 3μM deoxycytidine (dC) versus control media (n=3). **3g-j**. Relative viability dose response curve and calculated IC_50_ of MT3-KPC cells treated with **3g**. 5-aza-deoxycytidine, **3h**. 5-azacytidine, **3i**. Trifluorothymidine, or **3j**. Fialuridine in the presence of 3μM dC or control media (n=3). * *P* ≤ 0.05; ^**^ *P* ≤ 0.01; ^****^ *P* ≤ 0.0001

Pyrimidine-based chemotherapies represent a large and well characterized class of drugs (Fig. 3d) (Parker, 2009). Based on data in Fig.3b, c, the protective mechanism(s) of dC should be limited to those pyrimidine-based chemotherapies that share uptake or metabolism properties. To test this hypothesis, we first examined 5-fluorouracil (5-FU), a different class of pyrimidine nucleoside-based chemotherapy that is used for patients with PDA as a component of the combination chemotherapy FOLFIRINOX (Conroy et al., 2011). We found that the cytotoxic activity of 5-FU in PDA cells is not impacted by TEM CM or dC (Fig. 3e,f, Fig. S3h). Consistent with our model, the primary transporter utilized by 5-FU and the activating enzyme are distinct from those involved in dC uptake and metabolism (Table S3). We then tested a panel of other pyrimidine-based chemotherapies (Fig. 3g-j). We observed that the cytotoxic activity of 5-aza-deoxycytidine, but not 5-aza-cytidine, Fialuridine or Trifluorothymidine, was inhibited by concurrent treatment with 3μM dC (Fig. 3d). The transporters used by dC and the ribose-bearing chemotherapies tested share a high degree of overlap (Fig. S6c). Among the distinctions, intracellular activation by dCK differentiates anti-pyrimidine chemotherapies that are inhibited by dC from those that are not. The results from this chemical profiling strategy suggest that TEM-derived dC is inhibiting the cytotoxic activity of gem in PDA cells by reducing the effective concentration through molecular competition at dCK, the rate limiting enzyme for gem activation.

### Macrophages modulate gemcitabine response

We next sought to determine if macrophages in the tumor microenvironment in vivo contribute to gem resistance. To examine this, we utilized a syngeneic PDA tumor transplantation model in mice expressing a diphtheria toxin (DT) receptor under the Cd11b promoter (Zhang et al., 2017a), which allows for selective, temporary depletion of myeloid cells (Fig. 4a). Consistent with our hypothesis, we found the size of tumors treated with gem after depletion of myeloid cells to be decreased dramatically, as compared to single treatment or vehicle treated arms (Fig. 4b). The increased efficiency of gem in myeloid depleted tumors resulted in increased DNA damage, marked by γH2AX immunostaining (Fig 4c,d). In contrast, at end point, differences in proliferation or apoptosis were not evident among the groups (Fig. S4a). Further, consistent with our model on the role of TAM-derived dC, treatment with 5-FU-based chemotherapy in this model yielded no benefit from myeloid depletion (Fig. S4b).

**Figure 4:**
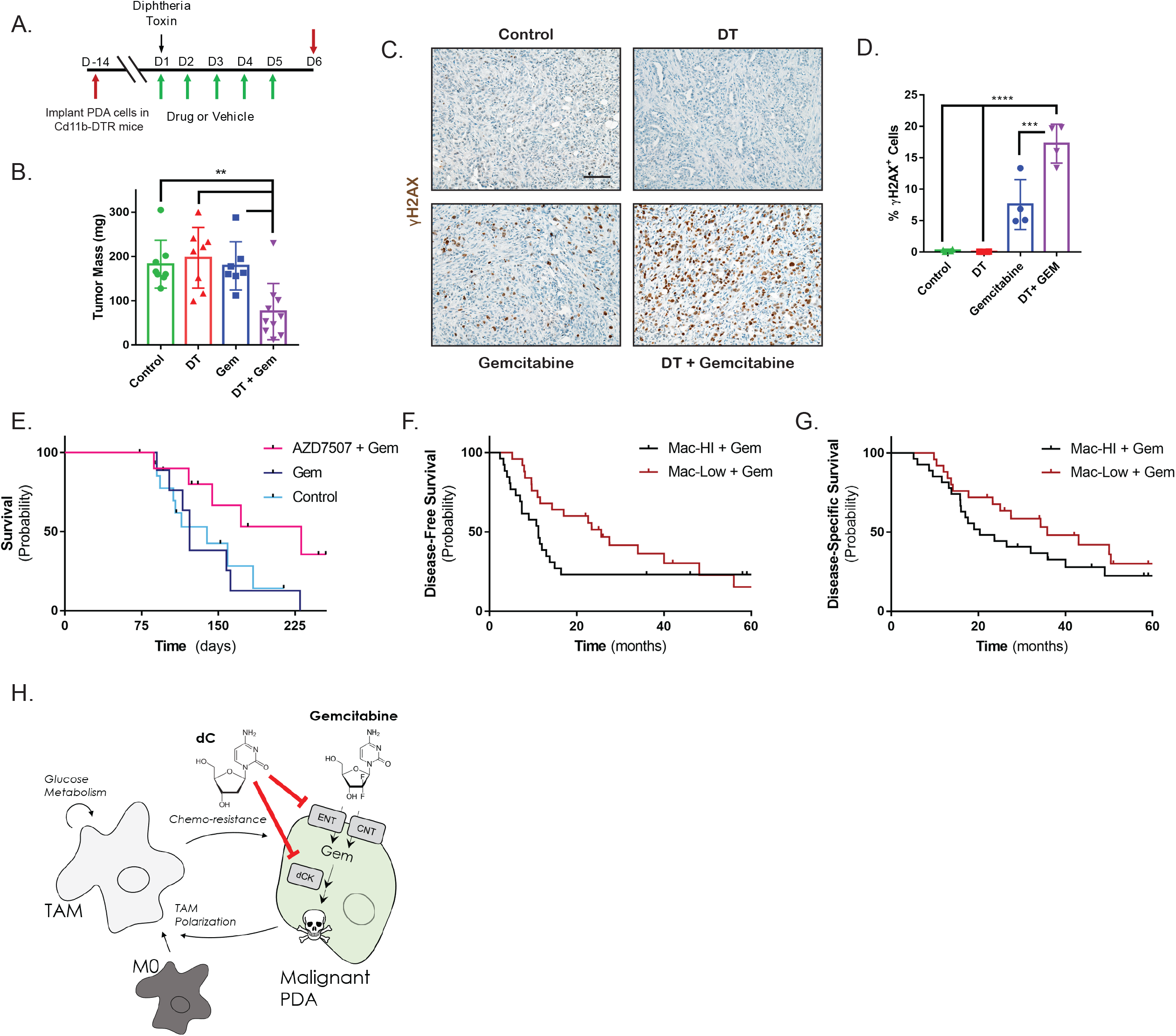
Macrophages inhibit gemcitabine treatment, burden predicts treatment response. **4a**. Schematic representation of Cd11b-DTR macrophage depletion tumor model with gemcitabine treatment schedule. **4b**. Mass of vehicle- (n=8), gemcitabine- (n=7), diphtheria toxin (DT)- (n=8), or DT + gemcitabine-treated (n=10) KPC-MT3 tumors at endpoint (day 6). **4c**. Immunohistochemical staining and **4d**. quantification of γH2AX in tumor tissue from (**4b**) (n=4). **4e**. Kaplan-Meier survival curve of control Kras^+/LSL-G12D^; Trp53^+/LSL-R172H^; Pdx1-Cre (KPC) mice (n=15), KPC mice treated with gemcitabine (n=9), or KPC mice treated with AZD7507 + gemcitabine (n=10). **4f**. Kaplan-Meier disease-free **4g**. or disease-specific survival of human PDA patients treated with adjuvant gemcitabine therapy with either high (n=26) or low macrophage (n=25) burden as determined by gene-expression signature. **4h**. Schematic representation of intratumoral PDA cell to TAM crosstalk. PDA cells program TAMs, which synthesize/release dC in a glucose metabolism-dependent manner. This inhibits the transport and activation of gemcitabine and thus gemcitabine cytotoxicity. ^**^ *P* ≤ 0.01; ^***^ *P* ≤ 0.001; ^****^ *P* ≤ 0.0001. Scale bar = 100μM.

To test the clinical benefit of targeting macrophages in combination with gem treatment, we treated a genetically engineered, autochthonous pancreatic cancer model (Kras^+/LSL-G12D^; Trp53^+/LSL-R172H^; Pdx1-Cre, known as KPC) with the Colony Stimulating Factor Receptor 1 (CSFR1) inhibitor AZD7507, which blocks the recruitment and activation of monocytes, in combination with gem. In this model, we observed prolonged survival compared to gem alone or control-treated mice (Fig. 4e). In a cohort of patients with PDA who underwent surgery and adjuvant gem, we also found significantly better survival in those with a low macrophage signature compared to those with a high macrophage signature (median disease-specific survival 35.8 months and 20.3 months, respectively) (Fig 4f,g). Additionally, macrophage high patients fare worse than macrophage low patients in absence of adjuvant therapy (Fig. S4c,d), indicating additional challenges which may be posed by macrophages, such as immunosuppression (Zhang et al., 2017a; Zhu et al., 2014).

In summary, our data demonstrate a previously undescribed mechanism of intra-tumoral metabolic crosstalk that promotes therapeutic resistance (Fig. 4h). In pancreatic tumors, malignant PDA cells recruit and polarize macrophages into TAMs (Liou et al., 2015; Zhang et al., 2017b). We found that TAMs impair the cytotoxic activity of the front-line chemotherapy gem through the release of dC. In the tumor, dC competes with gem based on their molecular similarity, and thereby reduces the therapeutic efficacy. Accordingly, since gem remains a backbone of the standard chemotherapy regimen for pancreatic cancer patients (Von Hoff et al., 2013), strategies to inhibit the recruitment of macrophages or reprogram anti-inflammatory macrophages have potential to increase the utility of this well tolerated and mainstay therapy, as has already been seen with FOLFIRONOX (Nywening et al., 2016).

## Discussion

Pyrimidine biosynthesis is energetically costly, suggesting that release is a regulated process. Purines are well-established signaling molecules (Di Virgilio and Adinolfi, 2017), and release of pyrimidine nucleosides at micromolar levels may suggest a similar function. Alternatively, metabolites and breakdown products of metabolism can act in a paracrine fashion to regulate metabolism, and these have important roles in physiology. Further, we and others have reported on examples of such processes among the cells in tumors (Lyssiotis and Kimmelman, 2017; Murray, 2016). We propose that the mechanism we discovered using a tumor model system is a previously undescribed physiological process, in which anti-inflammatory macrophages provide another cell type with pyrimidines during normal development. Indeed, a potential insight for the role of pyrimidines comes from a system-wide genetic deletion of the dC salvage enzyme dCK in mice. In these animals, development of lymphocytes is grossly impaired, demonstrating a requirement for pyrimidine salvage during lymphocyte development (Austin et al., 2012; Toy et al., 2010). Our data suggest that the source of the salvaged pyrimidines may be from macrophages. Furthermore, as TAMs release a panel of pyrimidine nucleosides, they may promote chemoresistance to other therapies.

Accordingly, these insights may shift the paradigm of how we consider the evolution of chemoresistance to nucleoside-like drugs, not only in PDA, but potentially in many other tumor types where TAMs play an important role.

## Acknowledgements

The authors would like to thank Drs. Luigi Franchi, Timothy Frankel, Filip Bednar, Vinee Purohit, and the Lyssiotis lab for critical discussion and scientific feedback and Daniel Long for histology support. CJH was supported by UL1TR000433, T32CA009676, and F32CA228328; LL by Pancreatic Cancer UK; SK T32GM113900; BN by T32CA009676 and T32DK094775; HCC by the Sky Foundation; JPM by Cancer Research UK; HSH by T32AI007413; CAL, HCC, and MPdM by a Cancer Center Support Grant (P30 CA046592) and U01 CA224145; MPdM by the American Cancer Society; CAL by the Pancreatic Cancer Action Network/AACR (13-70-25-LYSS), Damon Runyon Cancer Research Foundation (DFS-09-14), V Foundation for Cancer Research (V2016-009), Sidney Kimmel Foundation for Cancer Research (SKF-16-005), and the AACR (17-20-01-LYSS). Metabolomics studies were supported by DK097153, the Charles Woodson Research Fund and the UM Pediatric Brain Tumor Initiative.

## Author Contributions

CJH, MPdM, and CAL conceived of and designed this study. CJH and CAL planned and guided the research and wrote the manuscript. CJH, CP, HL, IK, LL, SD, DLK, PS, LZ, BN, HH, and SK performed experiments, analyzed, and interpreted data. DC, AB, HCC, JPM, MPdM, and CAL supervised the work carried out in this study.

## Materials and Methods

### Cell Culture

C57B6/J PDA lines KPC MT-3 and MT-5, were kind gifts from Dr. David Tuveson, KPC28258 from Dr. Sunil Hingorani, and KPC7940 from Dr. Gregory Beatty. iKras*3 PDA cells were previously described (Zhang et al., 2017a). MiaPaCa2, Panc1, BXPC3, L929, and RAW264.7 cell lines were purchased from ATCC. PA-TU-8902 was purchased from DSMZ. UM patient-derived cell lines were generated from de-identified patient tumor samples(Li et al., 2007), with the approval of the Institutional Review Board of the University of Michigan. All commercial cell lines were validated by STR profiling. All cells were maintained in high-glucose DMEM (Gibco) supplemented with 10% FBS (Corning), and routinely tested for mycoplasma contamination using MycoAlert (Lonza). iKras*3 PDA cells were additionally maintained in 1μg/mL doxycycline, and 1μg/mL doxycycline was added as a control to other conditions in experiments in which this cell line was used.

### Conditioned medium

Conditioned medium was produced by changing cell media of >50% confluent plates, then removing the medium from cells after 48 hours of growth and filtering through a 0.45μm polyethersulfone membrane (VWR). Boiled conditioned media was warmed to 100°C for 15 minutes and the precipitate was filtered out. Size cutoff columns (EMD Millipore, UFC900308) were used to remove species larger than 3kDa from the medium. The filtered conditioned medium was then resuspended in DMEM to the original volume. Conditioned media was used at a 3:1 ratio with fresh media to avoid effects of nutrient/metabolite exhaustion.

L929 conditioned media was prepared by incubating confluent L929 cells in fresh media for 48 hours, then filtered through a 0.45μm polyethersulfone membrane.

### Bone-marrow macrophage differentiation

Bone marrow was extracted from the femurs of C57B6/J mice as described (Celada et al., 1984), and cultured in macrophage differentiation media (i.e. high-glucose DMEM (Gibco) supplemented with 10% FBS (Corning), Sodium Pyruvate (Gibco), Penicillin/Streptomycin (Gibco), and 30% L929 conditioned media) for 5 days, refreshing on day 3.

### Macrophage polarization

Bone-marrow derived macrophages and RAW264.7 cells were polarized using either murine 10ng/mL M-CSF (Peprotech), 10ng/mL LPS (Enzo, ALX-581-011-L001), 10ng/mL murine IL-4 (Peprotech), or 75% PDA conditioned media. Macrophage subtypes were polarized from matched biological replicates.

### Metabolite sample preparation

Metabolite extraction from medium samples was done by adding 1 mL of conditioned medium to 4 mL of cold (−80°C) 100% methanol, then clarified by centrifugation. Intercellular metabolite fractions were prepared from cells that were lysed with cold (−80°C) 80% methanol, then clarified by centrifugation. Metabolite levels of intercellular fractions were normalized to the protein content of a parallel sample, and all samples were lyophilized via speed vac. Dried metabolite pellets from cells or media were re-suspended in 35 μL 50:50 MeOH: H_2_O mixture for metabolomics analysis.

^13^C-Glucose tracing was performed using glucose free DMEM (Gibco) supplemented with 10% dialyzed FBS (Gibco) and either 25mM ^12^C glucose (Sigma) or uniformly labeled ^13^C-glucose (Cambridge Isotopes). BMDM cultures were polarized with appropriate stimulus for 24 hours in normal media, then the media was replaced with ^13^C or ^12^C glucose labeling media and incubated overnight. Samples were then harvested and prepared as above.

### Metabolomics

For steady state metabolite profiling, an Agilent 1290 UHPLC-6490 Triple Quadrupole (QqQ) tandem mass spectrometer (MS/MS) system was used. For negative ion acquisition, a Waters Acquity UPLC BEH amide column (2.1 x 100mm, 1.7μm) was used with the mobile phase (A) consisting of 20 mM ammonium acetate, pH 9.6 in water, and mobile phase (B) acetonitrile. The following gradient was used: mobile phase (B) was held at 85% for 1 min, increased to 65% in 12 min, then to 40% in 15 min and held for 5 min before going to initial condition and held for 10 min. For positive ion acquisition, a Waters Acquity UPLC BEH TSS C18 column (2.1 x 100mm, 1.7μm) was used with mobile phase (A) consisting of 0.5 mM NH_4_F and 0.1% formic acid in water; mobile phase (B) consisting of 0.1% formic acid in acetonitrile. The following gradient was used: mobile phase (B) was held at 1% for 1.5 min, increased to 80% in 15 min, then to 99% in 17 min and held for 2 min before going to initial condition and held for 10 min. The column was kept at 40 °C and 3 μL of sample was injected into the LC-MS/MS with a flow rate of 0.2 mL/min. Tuning and calibration of the QqQ was achieved through Agilent ESI Low Concentration Tuning Mix.

Optimization was performed on the 6490 QqQ in negative or positive mode individually for each of 220 standard compounds to get the best fragment ion and other MS parameters for each standard. Retention time for each standard of the 220 standards was measured from pure standard solution or a mix standard solution. The LC-MS/MS method was created with dynamic (d)MRMs with RTs, RT windows and MRMs of all the 220 standard compounds.

In both acquisition modes, key parameters of AJS ESI were: Gas temp 275 °C, Gas Flow 14 L/min, Nebulizer at 20 psi, Sheath Gas Heater 250 °C, Sheath Gas Flow 11 L/min, Capillary 3000 V. For negative mode MS: Delta EMV was 350 V, Cycle Time 500 ms and Cell accelerator voltage was 4 V, whereas for positive acquisition mode MS: Delta EMV was set at 200 V with no change in cycle time and cell accelerator voltage.

The QqQ data were pre-processed with Agilent MassHunter Workstation Quantitative Analysis Software (B0700). Additional analyses were post-processed for further quality control in the programming language R. Each sample was normalized by the total intensity of all metabolites to reflect the same protein content as a normalization factor. Finally, each metabolite abundance level in each sample was divided by the median of all abundance levels across all samples for proper comparisons, statistical analyses, and visualizations among metabolites. The statistical significance test was done by a two-tailed t-test with a significance threshold level of 0.05.

For the ^13^C glucose tracing studies, an Agilent 1260 UHPLC combined with a 6520 Accurate-Mass Q-TOF LC/MS was utilized. Agilent MassHunter Workstation Software LC/MS Data Acquisition for 6200 series TOF/6500 series QTOF (B.06.01) was used for calibration and data acquisition. A Waters Acquity UPLC BEH amide column (2.1 x 100mm, 1.7μm) was used with mobile phase (A) consisting of 20 mM NH_4_OAc in water pH 9.6, and mobile phase (B) consisting of acetonitrile. The following gradient was used: mobile phase (B) was held at 85% for 1 min, increased to 65% in 12 min, then to 40% in 15 min and held for 5 min before going to initial condition and held for 10 min. The column was at 40 °C and 3 μL of sample was injected into the LC-MS with a flow rate of 0.2 mL/min. Calibration of TOF MS was achieved through Agilent ESI Low Concentration Tuning Mix.

Key parameters for both acquisition modes were: mass range 100-1200 da, Gas temp 350 °C, Fragmentor 150 V, Skimmer 65 v, Drying Gas 10 l/min, Nebulizer at 20 psi and Vcap 3500 V, Ref Nebulizer at 20 psi. For negative mode the reference ions were at 119.0363 and 980.01637 m/z whereas for positive acquisition mode, reference ions at 121.050873 and 959.9657 m/z

For ^13^C-labeling data analysis, we used Agilent MassHunter Workstation Software Profinder B.08.00 with Batch Targeted Feature Extraction and Batch Isotopologue Extraction and Qualitative Analysis B.07.00. Various parameter combinations, e.g. mass and RT tolerance, were used to find best peaks and signals by manual inspection. Key parameters were: mass tolerance = 20 or 10 ppm and RT tolerance = 1 or 0.5 min. Isotopologue ion thresholds and the anchor ion height threshold was set to 250 counts and the threshold of the sum of ion heights to 500 counts. Coelution correlation threshold was set to 0.3.

All other bioinformatics analyses including graphs and plots were done using R/Bioconductor.

### Chemicals

Gemcitabine, 5-azacytidine, decitabine, fialuridine, trifluorouridine, capecitabine, and 5-fluorouracil were sourced from Cayman Chemical. 6-Aminonicotinamide, 2-deoxyglucose, and all nucleosides and nucleotides were obtained from Sigma-Aldrich. Compounds were dissolved into PBS (Gibco) and added to media to treat cells. AZD7507 CSF1R has been previously described (Scott et al., 2013). FCCP, oligomycin, rotenone, antimycin A were from Sigma-Aldrich and stocks were prepared in DMSO; etomoxir, carnitine, palmitate, were from Sigma-Aldrich and prepared in water, BSA fraction V was from Roche and prepared in water.

### Cell Viability Assay

PDAC cells were grown on white walled 96-well plates (Costar 3917, Corning) at 1000 cells/well in triplicate and grown for four days. Cell viability was measured using the Cell Titer Glo 2.0 luminescence assay (Promega G9243). Luminescence was measured for 500ms using a SpectraMax M3 Microplate reader (Molecular Devices). ^IC^50 values were calculated using GraphPad Prism 7.

### Metabolic Flux Assay

To estimate the rate of glycolysis and mitochondrial respiration, a Seahorse Metabolic Flux Analyzer e96 XF instrument (Agilent) was used according to the manufacturer’s suggestion. 50,000 cells/well were seeded in the respective polarization culture media the day prior to the assay. The next day media was exchanged to the Seahorse assay media, containing 25 mM glucose, adjusted to pH~7.4. The cell plate was allowed to equilibrate for 1 hour in a non-CO_2_, 37°C incubator, followed by 3 sequential measurements for the basal respiration. The mitostress assay was performed by sequentially injecting 2μM oligomycin, 5μM FCCP, 0.5μM rotenone/0.5μM antimycin A. The cell number adjustment/normalization was performed using CyQuant NF (Thermo) after the assay.

### Seahorse Fatty Acid Oxidation Assay

Macrophages were polarized as described above. BSA-palmitate conjugates were prepared and the FAO assay was performed according to the manufacturer’s application notes with the following modifications: 50,000 cells/well supplied with the corresponding polarization factors in 5 replicates seeded the night before the assay. Incubation with the substrate-limiting media was omitted. The next day cells were washed twice with FAO media and incubated in for 1 hour in non-CO2 incubator. Etomoxir was added to final concentration 40μM to the corresponding control wells, cells were incubated for 15 mins, BSA-Palmitate or BSA were added immediately before the assay. The level of exogenous FAO was calculated as a difference between BSA-Palmitate and Etomoxir-Palmitate, and the respiration attributed to the proton leak was accounted for.

### Metabolic Pathway Analysis

Metabolic pathway enrichment analysis was performed using metabolites determined to be significant between M1 macrophages, M2 macrophages, and TEMs by two-tailed t-test with a significance threshold level of 0.05 using Metabolanalyst online software (http://www.metaboanalyst.ca/).

### RNAi

ON-TARGETplus siRNAs targeted toward murine Dhodh, Umps, or nontargeting sequence were purchased from Dharmacon. siRNAs were transfected into RAW246.7 cells using Lipofectamine RNAiMAX (ThermoFisher) per the manufacturer’s instructions. Media was changed to iKras*3 PDA conditioned media after 48 hours and allowed to condition for 48 additional hours. Media was then collected and filtered through a 0.45μM filter prior to use.

### Western blotting

Lysates were quantified by BCA assay (Thermo Fisher Scientific Inc., Waltham, MA) and equal protein amounts were run onto SDS-PAGE gels. Proteins were transferred from SDS-PAGE gel to Immobilon-FL PVDF membrane, blocked, then incubated with primary antibodies. After washing, membranes were then incubated in secondary antibody washed, then exposed on autoradiography film (Bioexpress) with West Pico ECL (Thermo Fisher Scientific).

### Antibodies

The following antibodies were used in this study, DHODH (Proteintech 14877), UMPS (Proteintech 14830), Vinculin (Cell Signaling 13901), Cleaved Caspase 3 (Cell Signaling 9664), Ki67 (Abcam ab15580), γH2A.X (Cell Signaling 9718), and Secondary anti-rabbit-HRP (Cell Signaling 7074).

^3^H-Gemcitabine DNA Incorporation Assay: PDAC cells were grown at near confluence for 12 hours in 6 to 60nM ^3^H-gemcitabine (14.8Ci/mmol) (Moravek Biochemicals, MT1572). DNA was harvested using a Qiagen DNeasy Blood and Tissue Kit and normalized according to concentration. The samples were diluted with 5mL of scintillation fluid and measured for 30 minutes using a Beckman LS6500 Scintillation Counter.

### Mouse strains

Kras^+/G12D^; Pdx1-Cre; Trp53^+/R172H^ (KPC) mice have been previously described (Hingorani et al., 2005). KPC animal experiments were performed under UK Home Office license and approved by the University of Glasgow animal welfare and ethical review committee. Mice were bred in house on a mixed background, maintained in conventional cages with access to standard diet and water ad libitum, and genotyped by Transnetyx (Cordoba, TN, USA). Mice of both sexes were randomly assigned to treatment cohorts at 10 weeks of age and treated with Gemcitabine (LC Labs) at 100mg/kg i.p. twice weekly +/-AZD7507 CSF1R inhibitor (AstraZeneca) at 100mg/kg p.o. twice daily; vehicle p.o. twice daily. Mice were culled at ethical endpoint by schedule 1 methods. CD11b-DTR experiments were performed as previously described (Zhang et al., 2017a), and conducted in accordance with the Office of Laboratory Animal Welfare and approved by the Institutional Animal Care and Use Committees of the University of Michigan. Gemcitabine was treated at 100mg/kg IP, Capecitabine at 500mg/kg PO.

### Histology

Mice were sacrificed by CO_2_ asphyxiation then tissue was quickly harvested and fixed overnight at room temperature with Z-fix solution (Anatech LTD). Tissues were processed using a Leica ASP300S Tissue Processor, paraffin embedded, and cut into 5 μm sections. Immunohistochemistry was performed on Discovery Ultra XT autostainer (Ventana Medical Systems Inc) and counterstained with hematoxylin. IHC slides were scanned on a Pannoramic SCAN slide scanner (Perkin Elmer), and annotation regions encompassing greater than 1mm of tissue were processed using Halo software (Indica Labs).

### Patient survival data

Patients were identified from the Australian Pancreatic Genome Initiative’s (APGI) contribution to the International Cancer Genome Consortium’s Pancreatic Cancer project (Dreyer et al., 2018). Patients with sufficient clinical data that underwent RNA sequencing (RNAseq) and analysis were included. All patients included underwent primary surgical resection for PDA and were defined as receiving Gemcitabine-based adjuvant therapy if completing 3 cycles or more. RNAseq and analysis was performed as previously described (Bailey et al., 2016). Macrophage infiltration signature was determined by gene enrichment analysis that defined upregulated gene expression associated with macrophage infiltration (Bailey et al., 2016; Rooney et al., 2015). Patients were dichotomized as high or low signature based on ranking the relative signature score from highest to lowest (Bailey et al., 2016). Clinical variables were determined using the AJCC 7^th^ staging system. Survival analysis was performed using the log-rank test using SPSS (Version 22.0; IBM SPSS Statistics, IBM Corporation, Armonk, NY). Deidentified patient data are provided in Supplemental Data Table 4.

### Statistical analysis

Statistics were performed using Graph Pad Prism 7 (Graph Pad Software Inc). Groups of 2 were analyzed with two-tailed students t test, groups greater than 2 with a single variable were compared using one-way ANOVA analysis with Tukey post hoc test, and groups greater than two multiple variables were compared with two-way AVONA with Tukey post hoc test. All error bars represent mean with standard deviation.

## Supplemental Figure Legends

**Supplemental Figure 1:** *TEMs derived from murine bone marrow provide gemcitabine resistance in a panel of PDA cell lines*. **S1a**. Schematic of macrophage differentiation and polarization paradigm. **S1b**. Representative relative viability gemcitabine dose response curves in the presence of 75% bone marrow-derived macrophage (BMDM) TEM conditioned media vs. control media. Error bars are s.d., each point a technical replicate (n=3). IC_50_ values derived from these dose response curves are plotted in **Fig. 1e**. **S1c**. Representative relative viability gemcitabine dose response curves in the presence of 75% RAW 264.7 (RAW) TEM conditioned media vs. control media. Error bars are s.d., each point a technical replicate (n=3). IC50s derived from these dose response curves are plotted in **Fig. 1f**. **S1d**. Representative relative viability gemcitabine dose response curves in the presence of complete DMEM supplemented with 3μM deoxycytidine (dC) vs. normal complete DMEM. Error bars are s.d., each point a technical replicate (n=3). IC_50_ values derived from these dose response curves were plotted in **Fig. 1m**.

**Supplemental Figure 2:** *Metabolic properties of macrophage subtypes.* **S2a,b** Seahorse mitostress assay of TEM, M1 and M2 macrophages measuring **S2a**. ECAR and **S2b**. OCR under standard conditions. Basal ECAR and OCR data were plotted in the bar graphs in **Fig.2b,c** and the Energy Profile in **Fig.2d**. Error bars are s.d., each point a technical replicate (n=5). **S2c**. Seahorse mitostress assay of TEM, M1 and M2 macrophages in fatty acid oxidation assay media supplemented with either Palmitate (Palm), Palm + etomoxir (ETO), or bovine serum albumin (BSA) + ETO as carbon source. Basal exogenous FAO data were plotted in the bar graph in **Fig.2e**. Error bars are s.d., each point a technical replicate (n=5). **S2d. I**sotopologue abundance of Itaconate from ^13^C-glucose in M1 vs. M2 macrophages. Error bars represent s.d. of biological replicates (n=3). **S2e.** Fractional isotopologue labeling from ^13^C-glucose in M1 vs. M2 macrophages. Error bars represent s.d. of biological replicates (n=3). **S2f**. Schematic summary of metabolic tracing data from uniformly labeled ^13^C-glucose in M1 vs. M2 macrophages. Black lines represent canonical glucose metabolism pathways. The red arrows represent metabolic pathway activity inferred from the tracing data in which M1 metabolic activity is greater than that of M2s; green arrows are used for metabolic pathways in which activity in M2s are dominant over that in M1s. The dashed lines indicate inferred pathway directionality based on labeling patterns. Acetyl-CoA, Acetyl coenzyme A; α-KG, alpha ketoglutarate; Asn, Asparagine; Asp, Aspartate; dC, deoxycytidine; Fum, Fumarate; Glu, Glutamate; Iso, isocitrate; Mal, Malate; OAA, Oxaloacetate; Suc, Succinate; UDP-GlcNAc, Uridine diphosphate N-acetyl-glucosamine; UDP-Glucose, Uridine diphosphate glucose.

**Supplemental Figure 3:** *Inhibition of de novo nucleotide biosynthesis blocks TEM mediated gemcitabine resistance.* **S3a**. Representative relative survival gemcitabine dose response curves for KPC-MT3 cells treated in normal glucose DMEM (25mM), 75% glucose restricted media (6.25mM), 75% conditioned media from TEMs grown in normal glucose DMEM, or 75% conditioned media from TEMs grown in glucose restricted media. Error bars are s.d., each point a technical replicate (n=3). IC_50_ values derived from these dose response curves are plotted in **Fig. 2l**. **S3b**. Representative relative survival gemcitabine dose response curves for KPC-MT3 cells treated in normal DMEM, normal DMEM + 200μM 2-deoxyglucose (2-DG), 75% TEM conditioned media, or 75% conditioned media from TEMs grown in 200μM 2-DG. Error bars are s.d., each point a technical replicate (n=3). IC_50_ values derived from these dose response curves are plotted in **Fig. 2m**. **S3c.** Representative relative survival gemcitabine curves for KPC-MT3 cells treated in normal DMEM, normal DMEM + 1μM 6-aminonicotinamide (6-AN), 75% TEM conditioned media, or 75% conditioned media from TEMs grown in 1μM 6-AN. Error bars are s.d., each point a technical replicate (n=3). IC_50_ values derived from these dose response curves are plotted in **Fig. 2n**. **S3d**. Relative viability of TEMs in PDA conditioned media alone, PDA conditioned media + 200μM 2-DG, or PDA conditioned media + 1μM 6-AN. Error bars are s.d., each point a technical replicate (n=3). **S3e**. Western blot analysis of gene knockdown of RAW 246.7 TEMs transfected with either nontarget siRNA, or siRNAs targeted to *Dhodh* or *Umps*. **S3f**. Representative relative survival gemcitabine dose response curves for KPC-MT3 cells treated in KPC-MT3 cells treated in normal DMEM, or 75% conditioned media from TEMs transfected with siRNA targeting *Dhodh*, *Umps*, or nontargeting (NT) siRNA. Error bars are s.d., each point a technical replicate (n=3). IC_50_ values derived from these dose response curves are plotted in **Fig. 2o**. **S3g**. Relative intra and extracellular abundance of gemcitabine in KPC-28258 cells or conditioned media, respectively, after 16 hours of treatment with 6nM gemcitabine in the presence or absence of 3μM deoxycytidine, as measured by LC-MS/MS. Error bars represent s.d., samples were prepared from individual plates (n=3). **S6h.** Representative relative viability dose response curve of MT3-KPC cells treated with 5-FU in the presence of 100μM deoxycytidine (dC) versus control media. Error bars represent s.d., each point represents a technical replicate (n=3). Significance was calculated by one-way ANOVA with Tukey post hoc test, ^**^ *P* ≤ 0.01; ^****^ *P* ≤ 0.0001.

**Supplemental Figure 4:** *Murine and human data for chemotherapy-treated pancreatic tumors.* **S4a**. Histology of KPC-MT3 tumors from vehicle-, gemcitabine-, diphtheria toxin (DT)-, or gemcitabine + DT-treated Cd11b-DTR mice. CC3, cleaved caspase 3; H&E, Hematoxylin and Eosin. **S4b**. Mass of KPC-MT3 tumors at endpoint from vehicle- (n=9), capecitabine- (n=10), DT-(n=8), or DT + capecitabine-treated (n=10) mice. Error bars represent s.d. **S4c.** Kaplan-Meier disease-free **S4d.** or disease-specific survival of human PDA patients with either high (n=15) or low macrophage (n=12) burden as determined by gene-expression signature. Lack of significance in **S4b** was determined by one-way ANOVA with Tukey post hoc test. Scale bar = 100μm.

## Supplemental Table Legends

**Supplemental Table 1**: Calculated abundance of pyrimidines in TEM conditioned media, determined via LC-MS/MS from 3 separate TEM preparations. SD = standard deviation.

**Supplemental Table 2**: Metaboanalyst pathway enrichment analysis.

**Supplemental Table 3**: Table of deoxycytidine and anti-pyrimidine metabolites, their reported transporters, activating kinases, and inhibition by deoxycytidine treatment. 5-aza-dC, 5-aza-dexoycytidine; 5-aza-C, 5-aza-cytidine; 5-FU, 5-fluorouracil; CNT, Concentrative nucleoside transporter; dCK = deoxycytidine kinase; ENT, Equilibrative nucleoside transporter; OAT2, organic anion transporter 2; TK, thymidine kinase; TYMS, thymidylate synthetase; UCK, uridine-cytidine kinase.

**Supplemental Table 4:** De-identified patient data.

## References

Austin, W.R., Armijo, A.L., Campbell, D.O., Singh, A.S., Hsieh, T., Nathanson, D., Herschman, H.R., Phelps, M.E., Witte, O.N., Czernin, J., et al. (2012). Nucleoside salvage pathway kinases regulate hematopoiesis by linking nucleotide metabolism with replication stress. J Exp Med 209, 2215–2228.

Bailey, P., Chang, D.K., Nones, K., Johns, A.L., Patch, A.M., Gingras, M.C., Miller, D.K., Christ, A.N., Bruxner, T.J., Quinn, M.C., et al. (2016). Genomic analyses identify molecular subtypes of pancreatic cancer. Nature 531, 47–52.

Celada, A., Gray, P.W., Rinderknecht, E., and Schreiber, R.D. (1984). Evidence for a Gamma-Interferon Receptor That Regulates Macrophage Tumoricidal Activity. Journal of Experimental Medicine 160, 55–74.

Conroy, T., Desseigne, F., Ychou, M., Bouche, O., Guimbaud, R., Becouarn, Y., Adenis, A., Raoul, J.L., Gourgou-Bourgade, S., de la Fouchardiere, C., et al. (2011). FOLFIRINOX versus gemcitabine for metastatic pancreatic cancer. N Engl J Med 364, 1817–1825.

Di Caro, G., Cortese, N., Castino, G.F., Grizzi, F., Gavazzi, F., Ridolfi, C., Capretti, G., Mineri, R., Todoric, J., Zerbi, A., et al. (2016). Dual prognostic significance of tumour-associated macrophages in human pancreatic adenocarcinoma treated or untreated with chemotherapy. Gut 65, 1710–1720.

Di Virgilio, F., and Adinolfi, E. (2017). Extracellular purines, purinergic receptors and tumor growth. Oncogene 36, 293–303.

Dreyer, S.B., Jamieson, N.B., Upstill-Goddard, R., Bailey, P.J., McKay, C.J., Biankin, A.V., and Chang, D.K. (2018). Defining the molecular pathology of pancreatic body and tail adenocarcinoma. The British journal of surgery 105, e183–e191.

Feig, C., Gopinathan, A., Neesse, A., Chan, D.S., Cook, N., and Tuveson, D.A. (2012). The pancreas cancer microenvironment. Clin Cancer Res 18, 4266–4276.

Halbrook, C.J., and Lyssiotis, C.A. (2017). Employing Metabolism to Improve the Diagnosis and Treatment of Pancreatic Cancer. Cancer Cell 31, 5–19.

Hingorani, S.R., Wang, L., Multani, A.S., Combs, C., Deramaudt, T.B., Hruban, R.H., Rustgi, A.K., Chang, S., and Tuveson, D.A. (2005). Trp53R172H and KrasG12D cooperate to promote chromosomal instability and widely metastatic pancreatic ductal adenocarcinoma in mice. Cancer Cell 7, 469–483.

Jha, A.K., Huang, S.C.C., Sergushichev, A., Lampropoulou, V., Ivanova, Y., Loginicheva, E., Chmielewski, K., Stewart, K.M., Ashall, J., Everts, B., et al. (2015). Network Integration of Parallel Metabolic and Transcriptional Data Reveals Metabolic Modules that Regulate Macrophage Polarization. Immunity 42, 419–430.

Koay, E.J., Truty, M.J., Cristini, V., Thomas, R.M., Chen, R., Chatterjee, D., Kang, Y., Bhosale, P.R., Tamm, E.P., Crane, C.H., et al. (2014). Transport properties of pancreatic cancer describe gemcitabine delivery and response. J Clin Invest 124, 1525–1536.

Li, C.W., Heidt, D.G., Dalerba, P., Burant, C.F., Zhang, L.J., Adsay, V., Wicha, M., Clarke, M.F., and Simeone, D.M. (2007). Identification of pancreatic cancer stem cells. Cancer Research 67, 1030–1037.

Liou, G.Y., Doppler, H., Necela, B., Edenfield, B., Zhang, L., Dawson, D.W., and Storz, P. (2015). Mutant KRAS-induced expression of ICAM-1 in pancreatic acinar cells causes attraction of macrophages to expedite the formation of precancerous lesions. Cancer Discov 5, 52–63.

Lyssiotis, C.A., and Kimmelman, A.C. (2017). Metabolic Interactions in the Tumor Microenvironment. Trends Cell Biol 27, 863–875.

Mitchem, J.B., Brennan, D.J., Knolhoff, B.L., Belt, B.A., Zhu, Y., Sanford, D.E., Belaygorod, L., Carpenter, D., Collins, L., Piwnica-Worms, D., et al. (2013). Targeting tumor-infiltrating macrophages decreases tumor-initiating cells, relieves immunosuppression, and improves chemotherapeutic responses. Cancer Res 73, 1128–1141.

Murray, P.J. (2016). Amino acid auxotrophy as a system of immunological control nodes. Nat Immunol 17, 132–139.

Nywening, T.M., Wang-Gillam, A., Sanford, D.E., Belt, B.A., Panni, R.Z., Cusworth, B.M., Toriola, A.T., Nieman, R.K., Worley, L.A., Yano, M., et al. (2016). Targeting tumour-associated macrophages with CCR2 inhibition in combination with FOLFIRINOX in patients with borderline resectable and locally advanced pancreatic cancer: a single-centre, open-label, dose-finding, non-randomised, phase 1b trial. Lancet Oncol 17, 651–662.

Olive, K.P., Jacobetz, M.A., Davidson, C.J., Gopinathan, A., McIntyre, D., Honess, D., Madhu, B., Goldgraben, M.A., Caldwell, M.E., Allard, D., et al. (2009). Inhibition of Hedgehog signaling enhances delivery of chemotherapy in a mouse model of pancreatic cancer. Science 324, 1457–1461.

Parker, W.B. (2009). Enzymology of Purine and Pyrimidine Antimetabolites Used in the Treatment of Cancer. Chem Rev 109, 2880–2893.

Perera, R.M., and Bardeesy, N. (2015). Pancreatic Cancer Metabolism: Breaking It Down to Build It Back Up. Cancer Discovery 5, 1247–1261.

Provenzano, P.P., Cuevas, C., Chang, A.E., Goel, V.K., Von Hoff, D.D., and Hingorani, S.R. (2012). Enzymatic targeting of the stroma ablates physical barriers to treatment of pancreatic ductal adenocarcinoma. Cancer Cell 21, 418–429.

Rhim, A.D., Oberstein, P.E., Thomas, D.H., Mirek, E.T., Palermo, C.F., Sastra, S.A., Dekleva, E.N., Saunders, T., Becerra, C.P., Tattersa, I.W., et al. (2014). Stromal Elements Act to Restrain, Rather Than Support, Pancreatic Ductal Adenocarcinoma. Cancer Cell 25, 735–747.

Rooney, M.S., Shukla, S.A., Wu, C.J., Getz, G., and Hacohen, N. (2015). Molecular and genetic properties of tumors associated with local immune cytolytic activity. Cell 160, 48–61.

Scott, D.A., Dakin, L.A., Daly, K., Del Valle, D.J., Diebold, R.B., Drew, L., Ezhuthachan, J., Gero, T.W., Ogoe, C.A., Omer, C.A., et al. (2013). Mitigation of cardiovascular toxicity in a series of CSF-1R inhibitors, and the identification of AZD7507. Bioorg Med Chem Lett 23, 4591–4596.

Siegel, R.L., Miller, K.D., and Jemal, A. (2016). Cancer statistics, 2016. CA Cancer J Clin 66, 7–30.

Sousa, C.M., Biancur, D.E., Wang, X., Halbrook, C.J., Sherman, M.H., Zhang, L., Kremer, D., Hwang, R.F., Witkiewicz, A.K., Ying, H., et al. (2016). Pancreatic stellate cells support tumour metabolism through autophagic alanine secretion. Nature 536, 479–483.

Toy, G., Austin, W.R., Liao, H.I., Cheng, D., Singh, A., Campbell, D.O., Ishikawa, T.O., Lehmann, L.W., Satyamurthy, N., Phelps, M.E., et al. (2010). Requirement for deoxycytidine kinase in T and B lymphocyte development. Proc Natl Acad Sci U S A 107, 5551–5556.

Van den Bossche, J., O’Neill, L.A., and Menon, D. (2017). Macrophage Immunometabolism: Where Are We (Going)? Trends Immunol 38, 395–406.

Vats, D., Mukundan, L., Odegaard, J.I., Zhang, L., Smith, K.L., Morel, C.R., Wagner, R.A., Greaves, D.R., Murray, P.J., and Chawla, A. (2006). Oxidative metabolism and PGC-1beta attenuate macrophage-mediated inflammation. Cell Metab 4, 13–24.

Von Hoff, D.D., Ervin, T., Arena, F.P., Chiorean, E.G., Infante, J., Moore, M., Seay, T., Tjulandin, S.A., Ma, W.W., Saleh, M.N., et al. (2013). Increased survival in pancreatic cancer with nab-paclitaxel plus gemcitabine. N Engl J Med 369, 1691–1703.

Weizman, N., Krelin, Y., Shabtay-Orbach, A., Amit, M., Binenbaum, Y., Wong, R.J., and Gil, Z. (2014). Macrophages mediate gemcitabine resistance of pancreatic adenocarcinoma by upregulating cytidine deaminase. Oncogene 33, 3812–3819.

Zhang, Y., Velez-Delgado, A., Mathew, E., Li, D., Mendez, F.M., Flannagan, K., Rhim, A.D., Simeone, D.M., Beatty, G.L., and Pasca di Magliano, M. (2017a). Myeloid cells are required for PD-1/PD-L1 checkpoint activation and the establishment of an immunosuppressive environment in pancreatic cancer. Gut 66, 124–136.

Zhang, Y., Yan, W., Mathew, E., Kane, K.T., Brannon, A., 3rd, Adoumie, M., Vinta, A., Crawford, H.C., and Pasca di Magliano, M. (2017b). Epithelial-Myeloid cell crosstalk regulates acinar cell plasticity and pancreatic remodeling in mice. Elife 6.

Zhu, Y., Knolhoff, B.L., Meyer, M.A., Nywening, T.M., West, B.L., Luo, J., Wang-Gillam, A., Goedegebuure, S.P., Linehan, D.C., and DeNardo, D.G. (2014). CSF1/CSF1R blockade reprograms tumor-infiltrating macrophages and improves response to T-cell checkpoint immunotherapy in pancreatic cancer models. Cancer Res 74, 5057–5069.

